# Single cell label-free probing of chromatin dynamics during B lymphocyte maturation

**DOI:** 10.1101/2021.01.12.426344

**Authors:** Rikke Morrish, Kevin Ho Wai Yim, Stefano Pagliara, Francesca Palombo, Richard Chahwan, Nicholas Stone

## Abstract

Large-scale intracellular signalling during developmental growth or in response to environmental alterations are largely orchestrated by chromatin within the cell nuclei. Chemical and conformational modifications of the chromatin architecture are critical steps in the regulation of differential gene expression and ultimately cell fate determination. Therefore, establishing chemical properties of the nucleus could provide key markers for phenotypic characterisation of cellular processes on a scale of individual cells.

Raman microscopy is a sensitive technique that is capable of probing single cell chemical composition - and sub-cellular regions - in a label-free optical manner. As such, it has great potential in both clinical and basic research. However, perceived limitations of Raman spectroscopy such as low signal intensity and the difficulty in linking alterations in vibrational signals directly with ensuing biological effects have hampered advances in the field. Here we use immune B lymphocyte development as a model to assess chromatin and transcriptional changes using confocal Raman microscopy in combination with microfluidic devices and correlative transcriptomics, thereby linking changes in chemical and structural properties to biological outcomes. Live B lymphocytes were assessed before and after maturation. Multivariate analysis was applied to distinguish cellular components within each cell. The spectral differences between non-activated and activated B lymphocytes were then identified, and their correlation with known intracellular biological changes were assessed in comparison to conventional RNA-seq analysis. Our data shows that spectral analysis provides a powerful tool to study gene activation that can complement conventional molecular biology techniques and opens the way for mapping the dynamics in the biochemical makeup of individual cells.

## Introduction

The ability to measure and quantify molecular changes during cellular development can enable the characterisation of cells during differentiation, cellular responses to extracellular cues, or disease progression. Conventional techniques, such as fluorescent tagging of molecules visualized with fluorescence microscopy(1) and transcriptomics and proteomics profiling (2), have been extensively used to assess molecular changes occurring within cells. However, their stark limitations are the need for target labelling and/or destruction of the biological specimens under study. That is why, non-invasive and label free vibrational spectroscopy techniques – including Fourier Transform Infrared (FTIR) and Raman microscopy – stand out.

Vibrational spectroscopy exploits the interaction between light and molecules to probe their vibrational modes in order to obtain a “chemical fingerprint” of a sample. Both FTIR and Raman have been used to monitor modifications to or changes in expression of specific biomolecules, such as DNA levels during cell cycle(3–7), protein modifications(8, 9) and DNA damage(10–13). However, it is becoming apparent that, although it is possible to identify specific signals associated with intracellular biochemical changes, a whole range of subtle spectral variations characterise cell state changes. This is not surprising, as cellular responses induce a swarm of transcriptional up- and down-regulation orchestrating changes to the transcriptomic and proteomic profile of the cell. Using multivariate analysis, spectral information enables classification of cell states or phenotypes of mammalian (14–25), bacterial, and yeast cells (26, 27).

It is this label-free classification that has great potential both in i) clinical settings, for disease diagnosis and prognosis, and ii) in biomedical research, for example in cell sorting for downstream processes. How powerful this tool can be, depends upon our understanding of the correlation between the spectral output and the underlying biochemical pathways within the cells. In bacterial research, antibiotic resistance is of great interest. Spectral markers of antibiotic resistance have been identified at the population level(26), and more recently a correlation between peak intensities and expression levels of antibiotic resistance contributing genes has been found (26). Importantly, this was done in the absence of antibiotics, indicating that the transcriptional profile of the given cells, affected on its turn by environmental changes(28), rather than their phenotypic response to the presence of antibiotics, were responsible for the spectral signatures (26). This correlation between Raman spectra and transcriptomic data has further been explored in a comprehensive manner in yeast where it has been shown that Raman spectra and transcriptomic data are linearly correlated (27). In both yeast and bacterial cells, several environmental conditions have been examined. A linear transformation matrix describing the relationship between the Raman data and the transcriptomic data has made it possible to predict an environment-specific Raman spectrum based on transcriptomic data. Conversely, the transcriptome of specific environment has been predicted based on the Raman data (27).

Transcriptomic readout consists of thousands of RNA transcripts, whereas Raman spectroscopy can measure the phenotypic expression of the RNA transcripts, i.e. the biochemical result of the transcriptomic profile. Noteworthy, all transcripts do not change independently; instead, strong correlations are found between transcripts that are controlled by global regulators, reflected in the Raman signals. In the yeast study, it has been determined that only 17 transcripts are sufficient for determining a linear correlation with the Raman spectra. The transcripts largely responsible for the linear correspondence have been identified by determining the variable importance in projection (VIP) values for each transcript. The top scoring transcripts were primarily non-coding RNAs in yeast and ribosome-related transcripts in bacteria (27). The correlation between them does not mean that the Raman spectra directly measure the expression levels of the transcripts in question. Instead, the downstream effects – i.e. changes to expression levels of large groups of genes and the resulting change in biochemical composition of the cells – are quantified by Raman. Thus, by analysing the correlation between Raman spectra and transcript expression levels, the key cellular pathways affecting the biochemical profile assessed by Raman spectroscopy may be identified.

To our knowledge i) the correlation between Raman spectra and transcriptomic readouts has never been studied in mammalian cells, and ii) it has not been examined in the context of cell differentiation. To explore this correlation in mammalian cells, B lymphocytes were chosen as a model cell system. Immune activation of these cells initiates large-scale changes to the transcriptomes, resulting in the differentiation of naïve B cells into mature B cells and class switch recombination (CSR) of the immunoglobulin receptor. Furthermore, as CSR requires reorganisation of the DNA, it is a highly regulated process. A large number of regulatory proteins and RNAs have been shown to be involved (29–35). However, the complex coordination of regulatory pathways and expression modulations is not yet fully understood. Novel techniques and approaches are needed to identify key regulatory RNAs and proteins previously unlinked to the B cell activation differentiation process.

## Materials and Methods

### CH12F3 cell culture and immune activation

CH12F3 cells were cultured in RPMI 1640 medium with 10% fetal bovine serum, 5% NCTC-109, 1% Pen-Strep, 1% glutamine, 1% sodium pyruvate and 50 μM β-mercaptoethanol. The cells were immune activated by incubating them with a cytokine cocktail (CIT) consisting of 2.5 μg/ml anti-CD40, 10 ng/ml IL-4 and 50 ng/ml TGFβ.

### Flow cytometry and antibodies

Flow cytometry measurements, for monitoring class switching assays of CH12F3 cells, were performed using a BD Accuri C6 Plus flow cytometer. 1% PFA fixed CH12F3 cells were stained with FITC Antimouse IgA antibody and APC Anti-mouse IgM antibody, both 1:200 dilution.

### Microfluidic device preparation

A microfluidic silicon mould was designed to create a small reservoir chip for maintaining cell viability during Raman measurements as previously described(36). This mould was then replicated in polydimethylsiloxane (PDMS) using a 9:1 ratio of base-to-curing agent. The PDMS was heated at 70°C for one hour and cut to size. A 1.5 mm biopsy punch was used to create an inlet and outlet within the reservoir. The chip was bonded to a glass coverslip using surface ionisation by oxygen plasma treatment (10 second exposure to 30 W plasma power in 1 mbar of air).

### Sample preparation and Raman mapping

Cells were washed in PBS, pelleted, and resuspended in PBS. PBS was flowed into the bonded chip using Portex tubing PE 0.86×0.33mm BxW using a syringe and a 21G microlance. Cells were flowed into the chip in the same way and left to settle for minimum 30 minutes. Raman maps were collected using a WITec alpha300R confocal Raman microscope system consisting of a 532 nm laser, a fibre-coupled UHTS spectrometer and an optical microscope with a 0.7 NA, 50× objective. The microfluidic chip containing the cells in PBS suspension was held coverslip side facing up in a custom holder. Single cells (adhering to the glass) were identified in white light imaging mode, then the focus was adjusted in Raman mode using the oscilloscope to maximise the scattered signal intensity (of the C-H stretching peak in the range 2700-3000 cm^-1^), and cells maps were collected with 5 measurement points per micrometre using a 0.1 s integration time per point. Cells were kept in the microfluidic chip for a maximum of four hours during measurements, before a new chip with fresh cells was prepared.

### Raman data analysis

Data processing was performed using MATLAB 2020a. Common k-means analysis with 10 clusters was performed on 118 cell maps (58×D0, 60×D4). This involved the calculation of similarity measures between each of the spectra from all 118 maps (a total of 716,250 spectra). As the most similar spectra are grouped together, the spectrum of the new group becomes the mean of its members. At the end of the process, when ten similar group clusters remained, they contained the spectra from regions of cells with similar biochemical constituents, each represented by a mean spectrum or centroid. Each of the 10 clusters was assigned as nucleus, cytoplasm or background, based on the spectral profiles of the cluster centroids. Hence, a mean nucleus, cytoplasm, whole cell (cytoplasm + nucleus) and background spectrum was extracted from each cell map. An array of 13 maps (5×D0, 8×D4) was discarded from the dataset before further analysis, since they either contained no pixels identified as nucleus or cytoplasm, or very few pixels associated with nucleus – with a nucleus size of less than 3 μm.

To remove the background signal (from the coverslip and PBS), the background spectra were subtracted from the nucleus and cytoplasm spectra for each map in three steps. (1) The spectra were baseline corrected by subtracting an offset (based on the mean intensity in the range 1780-1840 cm^-1^). (2) The background spectra were smoothed using a Savitzky-Golay filter (order = 2, framelength = 99) to reduce the effect of noise. (3) The smoothed background spectra were then subtracted from the nucleus, cytoplasm and whole cell spectra.

Principal Component Analysis (PCA) was performed on nucleus, cytoplasm and whole cell spectra. For each principal component, a t-test was used to determine if scores were significantly different between D0 and D4 cells. Linear Discriminant Analysis (LDA) was also performed to calculate a supervised classification model based on a combination of the PC scores. The resulting linear discriminant function could be used to single out the key peaks responsible for the discrimination between D0 and D4 cells.

### RNA extraction

Total cell RNA was extracted by TRIzol followed with chloroform for phase separation and 100% isopropanol for RNA precipitation. Total RNA was eluted in 30 μl RNase-free water after being washed twice in 75% ethanol. The RNA concentration was assessed using a NanoDrop 2000 spectrophotometer (Thermo Scientific, Waltham, MA, USA). The RNA yield and size distribution were analysed using an Agilent 2200 Tapestation with RNA Screentape (Agilent Technologies, Foster City, CA, USA).

### RNA-seq library preparation, next-generation sequencing and data processing

For small RNA library preparation, RNA aliquots were used for library preparation using NEBNext Multiplex Small RNA library preparation kit (New England Biolabs, Ipswich, MA, USA). The PCR amplified cDNA construct (from 140–160 bp) was purified using a QIAquick PCR Purification kit (Qiagen). The purified cDNA was directly sequenced using an Illumina MiSeq 2000 platform (Illumina, San Diego, CA, USA).

For long RNA library preparation, libraries were constructed using Ribo-Zero Magnetic Gold Kit (Human) (Illumina, San Diego, CA, USA) and NEBNext^®^ Ultra™ RNA Library Prep Kit for Illumina (New England Biolabs) according to the manufacturer’s instructions. Libraries were tested for quality and quantified using qPCR (Kapa Biosystems, Woburn, MA, USA). The resulting libraries were sequenced on a HiSeq 2500 instrument that generated paired-end reads of 100 nucleotides.

Raw sequencing reads were checked for potential sequencing issues and contaminants using FastQC. Adapter sequences, primers, number of fuzzy bases (Ns), and reads with quality scores below 30 were trimmed. Reads with a length of less than 20 bp after trimming were discarded. Clean reads were aligned to the mouse reference genome (GRCm38, 53,465 annotated genes in total) using the TopHat 2.0 program, and the resulting alignment files were reconstructed with Cufflinks (37). The transcriptome of each sample was assembled separately using Cufflinks 2.0 program.

### Sequencing data analyses and statistical methods

Read counts of each sample were subjected to cluster analysis (38) and differential expression analysis using RNA-seq 2G (39). Genes with |fold-change| ≥1, P value ≤0.05 and false discovery rate (FDR) ≤0.05 were considered statistically significant. Expression of significant differentially expressed genes in different B cell subsets was determined using My Geneset ImmGen (40). Interaction and gene targets of identified DE ncRNAs in cells and paired EVs were predicted by miRNet and ENCORI (41, 42). ncRNAs-target interaction network was constructed by Cytoscape v3.8.0 (43).

### Partial least squares regression

To analyse the potential correlation between the Raman data and the transcriptomic data, Partial Least Squares (PLS) regression was applied to the datasets. The transcriptomic data consisted of the read counts for 17,725 transcripts with three D0 samples (D0-1, D0-2, and D0-3) and three D4 samples (D4-1, D4-2, and D4-3). Dimension reduced Raman data were used in the form of PC scores. To correspond to the three replicates for each condition of the transcriptomic data, the Raman cell measurements were randomly assigned to three groups of D0 and three groups of D4.

PLS regression analysis was performed with a leave-one-out approach; each of the six samples was removed in turn. Each leave-one-out analysis determined a linear regression model between the Raman (R_-i_) and transcriptomic (T_-i_) datasets – meaning the PLS regression coefficients matrix, BETA_-i_ was found, so that

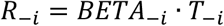

For each PLS regression analysis, Raman PC scores were predicted for the left-out sample, i, using the transcriptomic data (T_i_) and the BETA_-i_ matrix.

To assess the validity of the predicted Raman PC scores (and thus the regression model), they were compared against the single cell PCA scores. For further assessment, the predicted Raman PC scores were then converted to LDA scores and again compared with the single cell data.

## Results

### Identifying nucleus and cytoplasm in single cell Raman maps using common k-means

Raman maps were collected from 118 live CH12F3 cells suspended in isotonic PBS-filled microfluidic chambers. The cells remained in place throughout measurements. Common k-means analysis was applied to identify cell (vs background) pixels, as well as to distinguish the nucleus from the cytoplasm within each cell (Figure 1a-c). Additional examples are shown in Figure S1a-c. Inspection of the cluster centroid spectra (Figure 1b) informed the segmentation.

**Figure 1:**
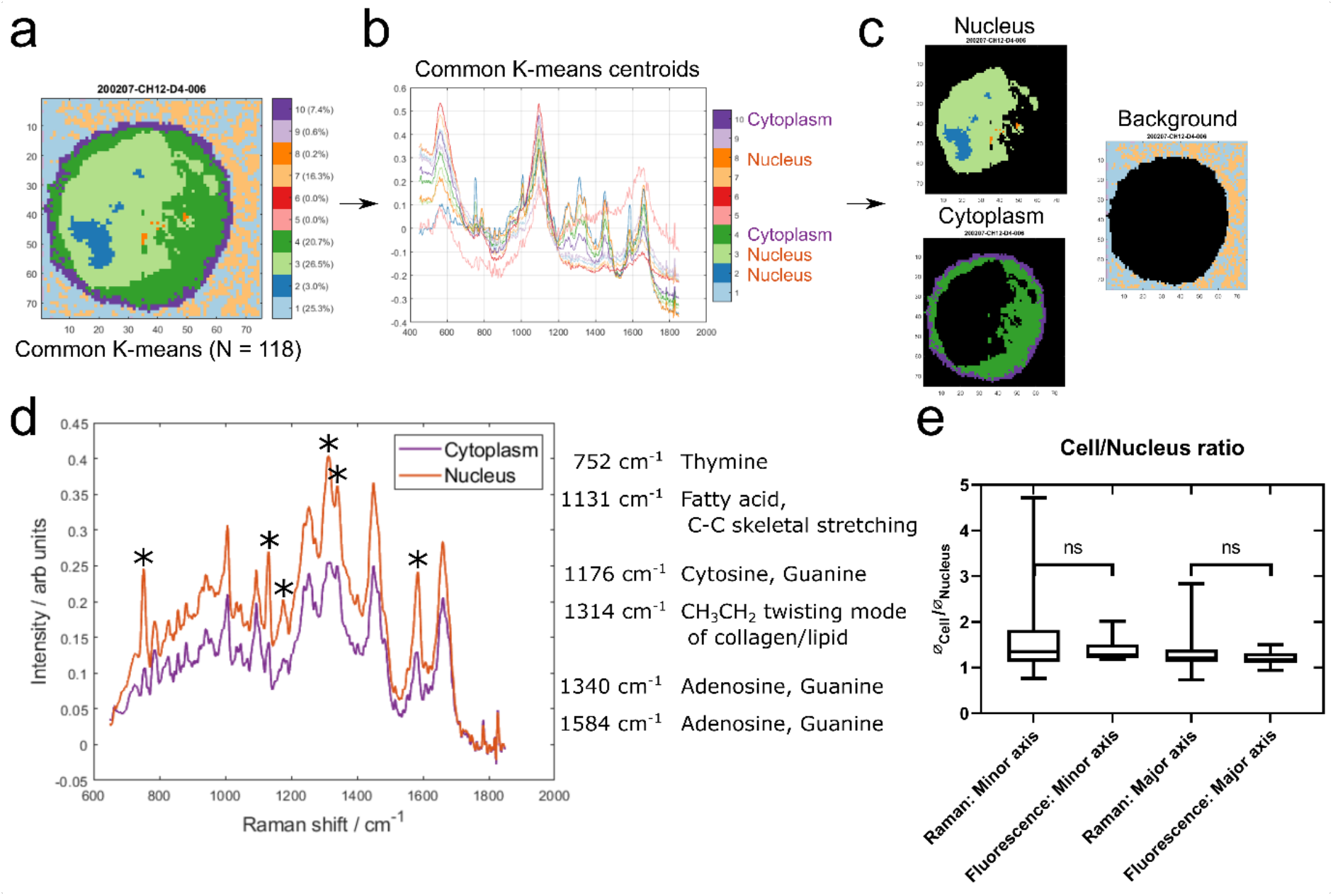
Identifying nucleus and cytoplasm associated areas using common K-means. **(a)** An example cell map (specifically sample 200207-CH12-D4-006) after common K-means with 10 clusters, which was used to analyse 118 individual cell maps concurrently. **(b)** The 10 common K-means centroid spectra. Spectral assignments were used to identify the clusters associated with cytoplasm (4 and 10) and nucleus (2, 3, and 8). **(c)** Example cell map (as seen in a) with nucleus (top left), cytoplasm (bottom left) and background (right) associated pixels highlighted. **(d)** Comparison of the mean cytoplasm and nucleus spectrum across all cells. The largest peak differences are highlighted and peak assignments are listed. **(e)** Comparison of nucleus/cell ratio between Raman maps and epifluorescence microscopy images.

The quality of the segmentation was assessed in two ways. Firstly, the mean nucleus spectrum was compared to the mean cytoplasm spectrum (Figure 1d). The most pronounced differences were associated with nucleic acid and lipid/fatty acid signals, with higher intensities found in the nucleus. Secondly, the size of the nucleus relative to the whole cell was assessed and compared with that from epifluorescence microscopy images of CH12F3 cells incubated with nucleic acid stains (Figure 1e). No statistically significant difference was found between the Raman and epifluorescence data. A larger variance was seen for the Raman data – possibly attributed to a number of smaller and kidney shaped nuclei (Figure S1f-g). These were not excluded as they were not outliers in the Raman spectral dataset.

### Quantifiable spectral differences between non-activated and activated B cells

To assess large-scale chromatin conformational and transcriptomic alterations, two groups of CH12F3 cells were compared: non-activated cells (D0) and cells at 96 hours post immune activation with cytokine (CIT: anti-CD40, IL-4 and TGFβ) cocktail (D4), as shown in Figure 2a. The immune activation of the cells was verified by quantifying the percentage of IgM-producing cells versus IgA-producing cells using flow cytometry (Figure S2).

**Figure 2:**
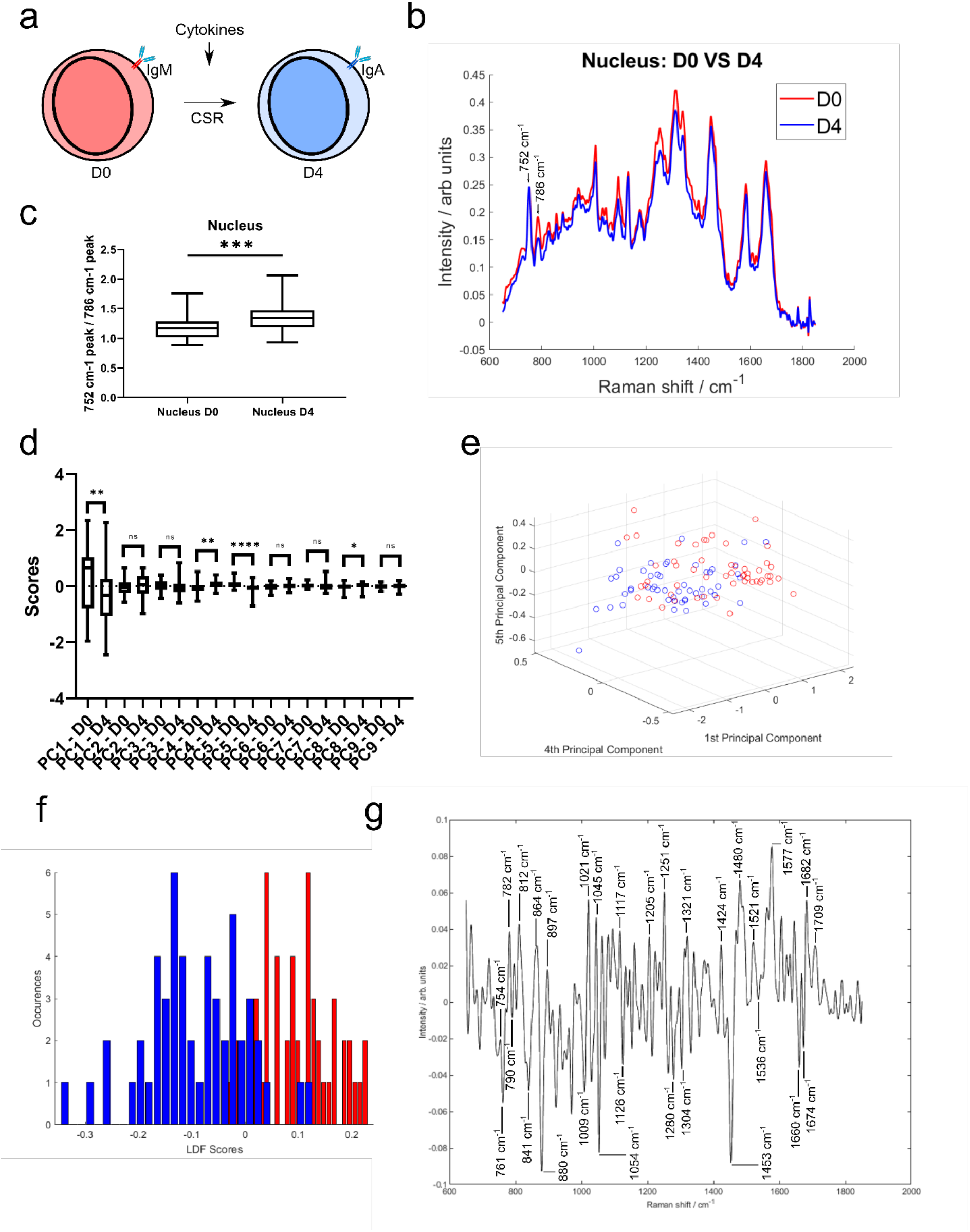
Quantifiable spectral differences between D0 and D4 cells. **(a)** Schematic representation of a CH12F3 cell undergoing class switch recombination in response to exposure to the cytokine cocktail. The expressed B cell receptor constant region changes from IgM to IgA. **(b)** The mean nucleus spectrum of D0 and D4 cells. Two neighbouring peaks are highlighted. **(c)** Peak ratio (752 cm^-1^ peak/786 cm^-1^ peak). A t-test gave a statistically significant difference between samples (ns.: P>0.05, *: <0.05, **: P<0.01, ***:P<0.001). **(d)** Principal Component analysis. Comparison of the first 9 PC scores. A t-test was applied to identify the principal components with a statistically significant difference between D0 and D4. **(e)** PCA analysis. Scores plotted for components 1, 4, and 5. **(f)** LDA analysis. Histogram of the distribution in the training model. **(g)** LDA analysis. Loading spectrum.

A prerequisite for further analysis and correlation with the transcriptomes was the ability to separate D0 and D4 cells based on their Raman spectra. Focusing on the nucleus, a number of spectral differences are apparent between D0 and D4 cells (Figure 2b). Namely, a peak at 786 cm^-1^ shows a large variation between the two activation states. The neighbouring peak at 752 cm^-1^ does not show this variation. Both of these peaks are associated with nucleic acids (44–46). The distribution of the 752 cm^-1^ to 786 cm^-1^ peak ratios was found to be significantly different between D0 and D4 cells (Figure 2c). This nucleic acid peak ratio therefore has the potential to provide a measure of activation status through the measurement of changes to DNA within the cells.

To further explore the spectral differences between D0 and D4 cells, multivariate approaches were applied. An unsupervised method, Principal Component Analysis (PCA), showed a separation between D0 and D4 cells (Figure 2d-e and Figure S3a-c). Four PCs had a statistically significant difference between D0 and D4 scores. The loading spectra of those showed a range of peaks associated with both nucleic acids, lipids and proteins (Figure S3d-g). Nucleic acid peaks around 786 cm^-1^ (PC1) and 752 cm^-1^ (PC1, PC4 and PC5) were amongst these, supporting the use of that peak ratio to distinguish between D0 and D4. Although a number of other nucleic acid peaks were identified, it is clear that intracellular changes of protein and lipid are also drivers for the spectral differences.

A supervised method, Linear Discriminant Analysis (LDA) was then applied, building on the PC scores and determining a classifier to discriminate between D0 and D4 cells. The two groups showed a very good separation (Figure 2f). Using a leave-one-out analysis, it was determined that the LDA classifier had a sensitivity of 73.1% and a specificity of 81.1% for identification of D4 cells. The loading plot for the classifier, representing the spectral data separating D0 and D4 cells, consisted of a range of peaks (Figure 2g). Nucleic acid peaks in the 751-790 cm^-1^ range are again present. The largest peaks include nucleic acid, protein, sugar and lipid, such as guanine and cytosine (782 cm^-1^, 1251 cm^-1^, 1577 cm^-1^(47, 48)), phosphodiester (812 cm^-1^, 897 cm^-1^, 1424 cm^-1^(47)), tryptophan (754 cm^-1^, 761 cm^-1^, 880 cm^-1^(48–52)), polysaccharide structure and glucose (841 cm^-1^, 1117 cm^-1^(46, 53, 54)) and CH_2_ deformation (1304 cm^-1^, 1321 cm^-1^(46, 55)).

These results show that is possible to distinguish between D0 and D4 cells based on their Raman spectra. Peaks associated with nucleic acids are important for this separation, but alterations to other biomolecules are also detectable.

### Differing transcriptomic profiles of non-activated and activated B cells

The transcriptomic profiles of D0 and D4 cells were determined and analysed. Read counts were measured for a total of 17,725 transcripts, and differential gene expression analysis using DESeq2 was applied to identify genes that were up-or down-regulated in response to immune activation. Figure 3a shows transcripts with the largest change of expression between D0 and D4 samples clustered based on Euclidean distance. The two transcripts *Ighm* and *Igha*, which code for the immunoglobulin heavy chain constant regions of IgM and IgA respectively, are highlighted. *Ighm* expression is down-regulated in D4, while *Igha* is up-regulated. This is a hallmark of the CH12F3 class switching response and in agreement with the IgM to IgA isotype switching measured by flow cytometry (Figure S2).

**Figure 3:**
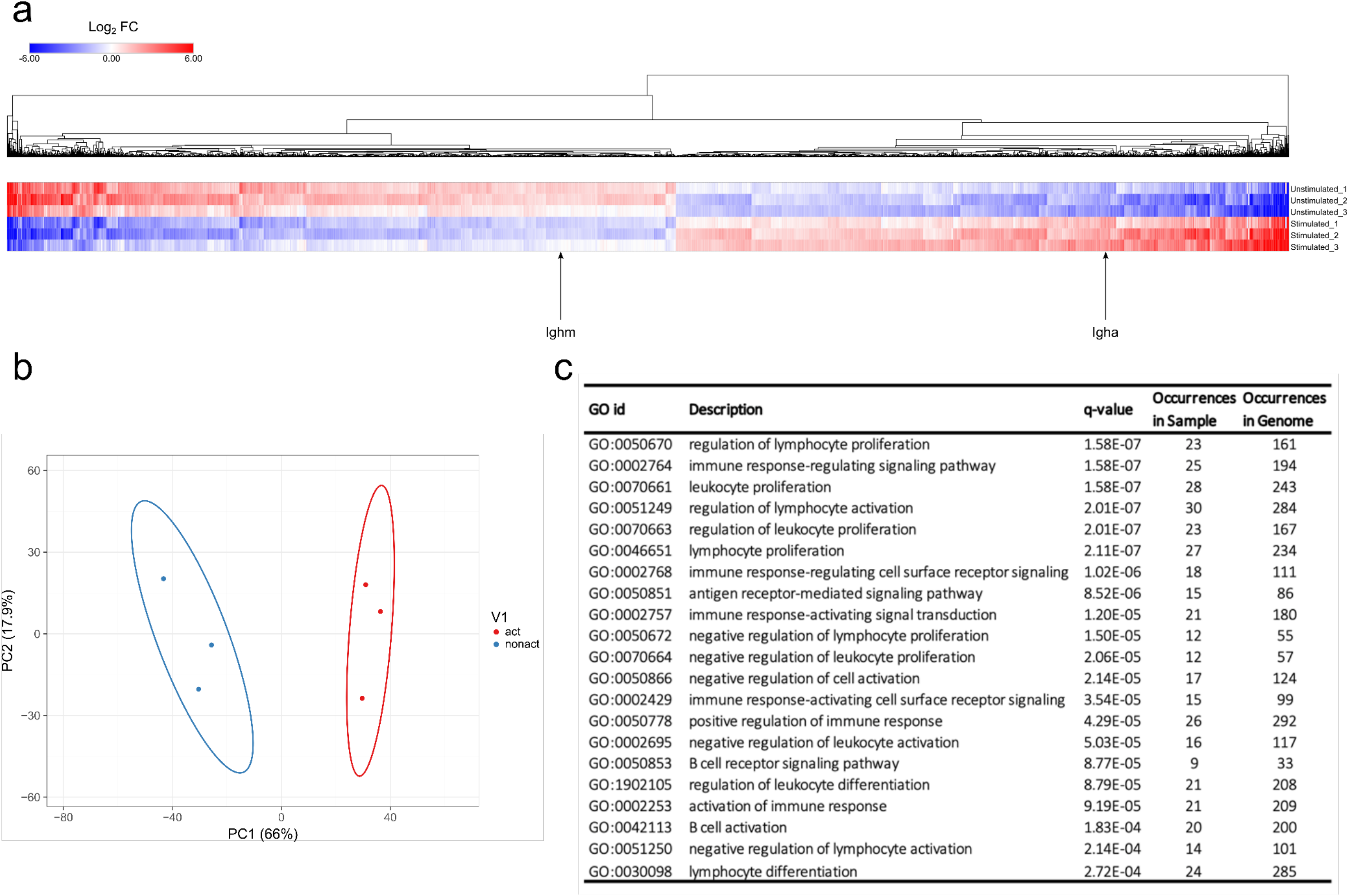
Transcriptomes of D0 and D4 cells. **(a)** Heatmap of the log2 fold change of transcripts from D0 and D4 samples, calculated using the DESeq2 software. Transcripts with |log2 fold change|>0.5 and FDR<0.05 are shown. Two transcripts, *Ighm* and *Igha*, are highlighted. *Ighm* and *Igha* code for the immunoglobulin heavy chain constant regions of IgM and IgA receptors, respectively. **(b)** PCA analysis showing separation of D0 and D4 samples. **(c)** Functional enrichment analysis of differentially expressed transcripts between D0 and D4. Table displaying significantly enriched gene ontology (GO) terms and associated biological processes.

The expression profiles of D0 and D4 cell samples clearly illustrate that several thousands of genes are up-or down-regulated upon immune activation of CH12F3 cells. These data are in line with published results regarding genes that are over-expressed during B cell maturation(56–59). For example, we could identify that genes AID, Bcl11a, CD40, and Ccr6 go up 2-, 3-, 2.6-, 8.7-fold in D4 compared to D0, respectively. This further illustrates the validity of our RNA-seq results. Further whole transcriptomic profiling, including PCA analysis (Figure 3b) and network comparison (Figure 3c), demonstrated a clear separation between D0 and D4 samples and the validity of our experimental approach.

PCA analysis on the expression profiles of D0 and D4 CH12F3 cohort samples were performed using Clustvis software tool(38). Differences were assessed after log2 transformation of normalised read counts at a threshold of *p* <0.05 for multiple comparisons. The variation of expression profile between D0 and D4 CH12F3 was displayed in first and second dimensions (PC1 vs PC2). Statistical significance was set at false discovery rate (FDR)<0.05. As a result, we could observe that the three D0 samples cluster in proximity together, as do the three D4 samples. But the D0 versus D4 clustered markedly separately from each other when plotted on the same graph (Figure 3b). This further validates the distinction in overall transcriptional profile of our cohorts. To further identify the specific basis for this distinction, we used Genemania pathway analysis tool(60). After including all hits in D4 expressed at log_2_ fold change > 1 with FDR < 0.05, we identified the top pathways to include the chemokine signalling pathway, leukocyte activation, immune cell differentiation, and B cell activation amongst other B cell related processes (Figure 3c). This further validates the specificity of our experimental design and its consistency.

### Linear correlation between transcriptomic and Raman data

Alterations in gene expression can ultimately cause changes in intracellular protein levels, as well as in other biomolecules through changes to metabolic pathways and intracellular structures. All these changes are bound to affect Raman spectral readouts. That a correlation exists between transcriptomic data and Raman spectra, as demonstrated for yeast and bacteria(27), is therefore not unexpected, albeit it was hard to predict whether a linear correlation would exist in a complex mammalian cell such as a B lymphocyte.

To test this hypothesis, a Partial Least Squares (PLS) regression analysis was applied to create a model for the prediction of Raman data from transcriptomic data of CH12F3 cells. A PLS regression model was determined from three D0 samples (D0-1, D0-2, D0-3) and three D4 samples (D4-1, D4-2, D4-3) of transcriptomic and Raman data (Figure 4a and Figure S4a). Using a leave-one-out approach, the validity of the linear regression model was tested on each sample in turn; one sample, i, was left out, and a PLS regression coefficients matrix, BETA_-i_, was determined from the remaining five samples. This matrix was then used to predict the Raman scores of the left-out sample.

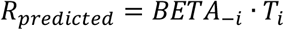

**Figure 4:**
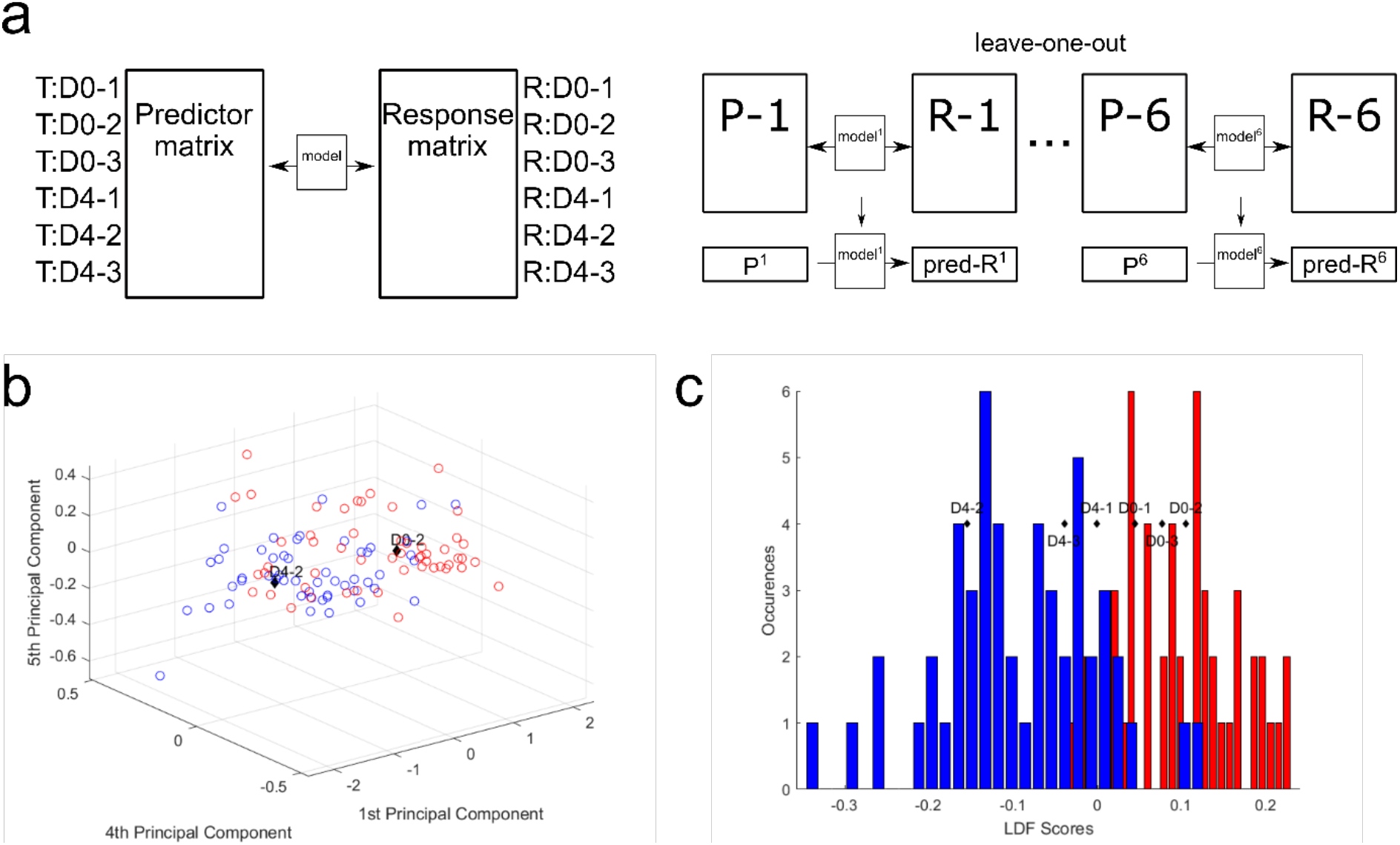
Partial least squares regression model correlates Raman and transcriptomic data. **(a)** Partial least squares regression model. **(b)** Raman scores predicted from transcriptomic read counts. Predicted D0-2 and D4-2 plotted with the single cell scores from PC1, PC4, and PC5. **(c)** Predicted Raman scores converted to LDA scores and plotted against the LDA scores histogram.

To assess the validity of these predictions and thus the PLS regression model, the predicted Raman data were compared to the single cell D0 and D4 data for each sample. Plotting the predicted PC scores against the single cell scores shows D0-2 and D4-2 within their expected regions (Figure 4b) and the rest around the intersection between D0 and D4 (Figure S4b-e). Further, by converting the predicted PC scores into their respective LD scores, the predicted group membership (D0 or D4) could be assessed (Figure 4c). The three D0 samples are found within the D0 region, while the D4 samples are found within the D4 region.

These results show that a linear correlation exists between transcriptomic profiles and Raman spectra of CH12F3 cells. Specifically, the variation in transcript expression levels between D0 and D4 cells is reflected in variation in Raman spectra of D0 and D4 cells – and transcriptomic data can be used to predict Raman data of CH12F3 cells.

### Identification of key transcripts for the correlation between Raman data and transcriptomic data

The importance of each transcript for the regression model is of particular interest, as this may reveal genes or pathways that are essential for the immune activation process. As shown in Figure 3, thousands of transcripts are significantly differentially expressed between D0 and D4. However, translation levels, protein modifications, and other regulatory mechanisms add further complexity to the final biochemical composition of the cell. A transcriptional profile does not account for these additional layers of regulation. Identifying transcripts of high importance for the correlation with the intracellular biochemical changes as measured by Raman microscopy may therefore be of great value. The Variable Importance in Projection (VIP) score was determined for each transcript. The top 20 transcripts for the PLS regression are shown in Figure 5a. We term these hits the VIP list. Upon further analyses of the protein coding entries in our VIP list, we could identify that many of our identified hits do indeed correlate with expression profiles from *in vivo* activated B cells isolated from murine germinal centre splenocytes (Figure 5b). Moreover, their expression quantifications (Figure 5c) correlate with post-activation B cell responses. It is worth noting that germinal centre splenocytes and CH12F3 cells are not directly comparable given the immortalised nature of the CH12F3 cell line. That is why we configured the *in vivo* germinal centre B cell response into three broad groups that we termed pre-, mid-, and post-activation (Figure 5b-c). We also took two representative *in vivo* cohorts for each of these three broad groups as represented in Figure 5b to ensure maximum congruency between *ex vivo* and *in vivo* analyses.

**Figure 5:**
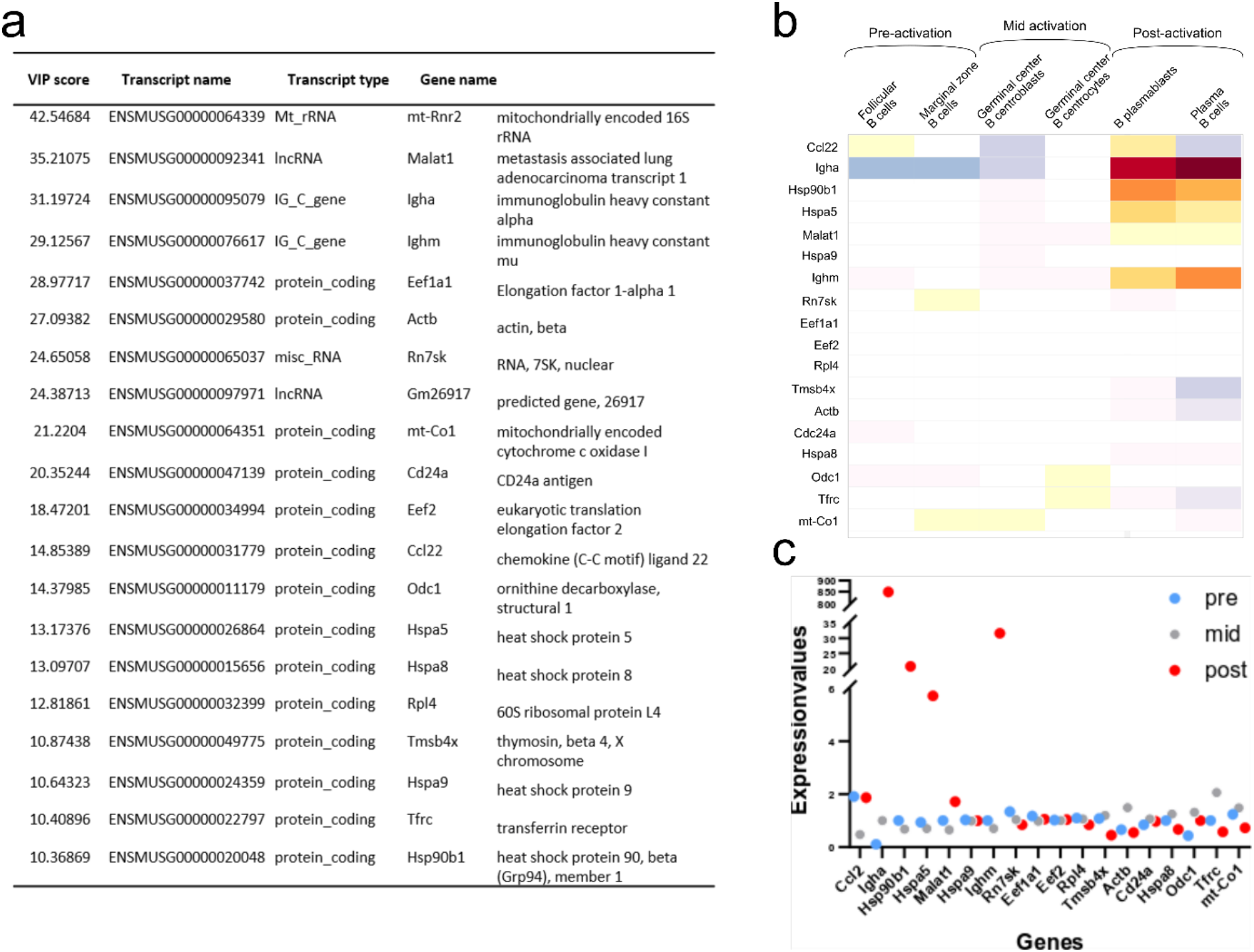
VIP transcript list and their expression *in vivo*. **(a)** The 20 transcripts with the highest Variable Importance in Projection (VIP) scores. **(b)** Expression of VIP genes *in vivo*. Median normalised gene expression values in different B cell subsets *in vivo* as annotated in Immunological Genome Projects. Further subclassified into pre-, mid-, and postactivation correlating to activation for CSR in CH12F3. **(c)** Quantification of normalised expression values of VIP genes in the dot plot.

*Ighm* and *Igha* are both found in the top four gene hits of the VIP list (Figure 5a). Although their change in expression levels results in the isotype switching from IgM to IgA, it is worth noting that they by no means are the most differentially expressed genes (Figure 3). Their high importance for the correlation with the Raman data therefore underlines that the transcripts with the highest fold change are not necessarily the most informative of the changing biochemical composition of a cell. The importance of IgM and IgA expression in the immune activation process is obvious, and their high presence on the VIP list supports the validity of the PLS regression model. Additional transcripts in the top 20 include regulatory and ribosomal RNAs, which is in line with data from yeast and bacterial analysis(27). A large number of regulatory proteins are also on the list, including a number of heat shock proteins, which have previously been shown to be important for CSR(61). Actin is also high on the list, in agreement with studies showing a regulatory role of the actin cytoskeleton in B cell activation (62, 63), as well as possibly the role of monomeric actin in DNA damage response (DDR) and chromatin modifications which happens during cell development or DNA repair(64–66).

A PLS regression model was also determined for whole cell and cytoplasm Raman data (Figure S5a-b). These allowed for predictions in line with the nucleus data (cytoplasm shown in Figure S5c-h). The top 20 VIP transcripts for those models (Table S1 and Table S2) were largely identical to the nucleus list. The values and order of the transcripts varied slightly, but only one transcript differed between the nucleus and whole cell list. For cytoplasm compared to nucleus, only three transcripts were different in the top 20 hits.

## Discussion

The cell segmentation used to isolate nucleus and cytoplasm regions within each cell using the Raman spectral maps proved useful for highlighting DNA Raman peaks. The 752/786 cm^-1^ peak ratio, shown to be statistically different between D0 and D4, has potential as a measure of activation status. If it is to be used as such, the biological significance of this peak ratio is of interest. The structure of DNA likely plays a role here. There are three biologically relevant double helical structures of DNA: A-DNA, B-DNA, and Z-DNA. B-DNA is the most common. A Raman peak at ~784-787 cm-1 has been shown to have a strong intensity for B-DNA, but much lower for the other two. The peak consists of two subpeaks; the breathing mode of the cytosine ring and the phosphodiester symmetric stretch of B-DNA backbone(67, 68). During B to Z transition of DNA, the phosphodiester symmetric stretch signal downshifts(68). As Z-DNA is associated with the rate of transcription(69, 70), it is plausible that the restructuring of DNA during activation could account for the difference between D0 and D4.

Looking at the whole Raman spectrum and downstream analyses, it is apparent that the spectra identified as from the nucleus are unlikely to be “pure” spectra (completely free of cytoplasmic signal). The similar VIP transcript lists of especially ‘nucleus’ and ‘whole cell’ analysis support this. Although a confocal microscope was used, optical signal from cytoplasm above and below the nucleus was likely measured too. The relatively large nucleus in CH12F3 cells and the round shape of the cells could have contributed to this. For larger and flatter adherent cells with smaller nuclei relative to the overall cell size, this may not occur to the same extent. Here it could be interesting to determine if a PLS regression model and its top VIP transcripts differed more between distinct cellular regions than for CH12F3 cells.

Both the PCA and LDA analysis revealed a myriad of spectral differences that allowed for the classification of D0 versus D4 cells. These included a large number of nucleic acid associated peaks, but also protein, lipid and sugar peaks. Classification of cell types or cell states based on Raman spectra has great clinical and research potential. However, understanding the biological significance of the spectral changes is of importance if these tools are to be implemented as a standard technique in biological laboratories. Peak assignments based on single molecule measurements provide some help with interpretation of the spectral changes. Correlation with transcriptomic data and identification of top VIP transcripts could add further value to the Raman data.

Here we showed that a linear correlation between Raman data and conventional next generation transcriptomic data exists in CH12F3 cells. Raman data were predicted based on transcriptomic profiles. When comparing the predicted Raman data to single cell data, the classification of each prediction was within the expected groups (D0 vs D4). We also identified the transcripts with the highest importance for the correlation with Raman spectra of non-activated and activated CH12F3 cells (Figure 5a). The immunoglobulin genes *Ighm* and *Igha* both featured in our top hits, highlighting the value of the PLS regression model as a valid phenotypic measurement for B cell activation. A number of regulatory RNAs and proteins were also in the top 20, some known to be involved in the regulation of CSR and activation, and others not previously shown to be involved. Further experiments exploring the role of these transcripts in B cell activation and CSR could be of great interest in the field of adaptive humoral immunity. This also suggests that our methodology could have the potential to identify novel molecular factors that other conventional assays might miss. One possible reason is the ability of our assay to combine both qualitative and quantitative analysis of nuclear signals along with a phenotypic readout of the overall status of the nucleus at the single cell level. RNA-seq, on the other hand, primarily measures quantitative readouts of bulk cells. Supported by similar results previously achieved in yeast cells(27), our work could provide a compelling argument for the use of Raman microscopy for phenotypic screening of a range of complex cellular processes.

Additional time points between D0 and D4 could further elucidate the correlation between Raman spectra and transcriptomic profiles. As there is some, although minor, inter-sample variability due to confluency levels and number of cell passages, it may also be beneficial to extract RNA and measure Raman spectra of cells from the same population on the same day. It would also be very interesting in the future to combine single-cell transcriptomics with Raman measurements. This would be a very powerful approach to unravelling the direct relationship between the two complementary data types. Indeed, there is still much to be determined, but there most certainly is a correlation between transcriptomic profiles and Raman spectra and this correlation can be further enhanced by combining more refined biological techniques. Understanding the origin of this correlation in different case studies will add value to Raman measurements of biological samples and aid interpretation of spectral changes. Furthermore, the work outlined here, suggests that Raman may also aid in the identification of key regulatory transcripts for immune activation. Future work will demonstrate if this can be translated to elucidating other cellular developmental processes, occurring in healthy cells or during disease.

## Data availability

The raw data supporting the conclusions of this manuscript will be made available by the authors, without undue reservation, to any qualified researcher.

## Author contributions

FP, RC, SP, and NS conceived and designed the experiments. RM and SP constructed the microfluidic chip. RM performed the experiments. KHWY acquired the transcriptomic data. RM, KHWY, and NS analyzed the data. RM wrote the manuscript with inputs from the other authors.

## Supplementary Material

The Supplementary Material for this article can be found online at:…

## Supplementary data

### Assessing the cell and nucleus segmentation

The additional cell map examples (Figure S1a-c) provide a clearer idea of how the cell and nucleus segmentation worked. Three clusters were determined to be associated with nucleus. The primary nucleus cluster varies between cell maps. That is also the case for the cytoplasm and background clusters. The Hierarchical Clustering Analysis (HCA) plot (Figure S1d) depicts the relationship between the ten clusters. All three nucleus clusters (2, 3, and 8) are similar to each other. One cytoplasm cluster (4) resembles the nucleus clusters, while the other one (10) is more similar to the background clusters. The ten clusters were assigned to either nucleus, cytoplasm or background based on the centroid spectra (Figure 1b), the HCA plot, and their effect on the cell and nucleus segmentation within each cell map. The cell versus background segmentation was straightforward, as seen in both the example maps and the centroid spectra. Nucleus versus cytoplasm segmentation was based on peaks associated with nucleic acid, which is more abundant in the nucleus. Cluster 4 proved to be the most difficult to assign – it was found to be at the interface between nucleus and cytoplasm, as shown by the example maps and HCA plot. The assignment of cluster 4 to cytoplasm was based on the centroid spectrum and the resulting nucleus size distribution of all maps.

The size and the shape of the nuclei vary between maps, as seen in Figure 1c and Figure S1a-c. This variation was quantified and assessed by measuring the major and minor axes of each nucleus and comparing them to those derived from epifluorescence images (Figure S1e-g). For both major and minor axes, the Raman mapped nuclei were smaller than the epifluorescence images nuclei. As both the measurement techniques and experimental setup were different between the two, this was not of great concern. Indeed, the Cell/Nucleus ratio was similar between the Raman maps and epifluorescence images (Figure 1e).

**Figure S1:**
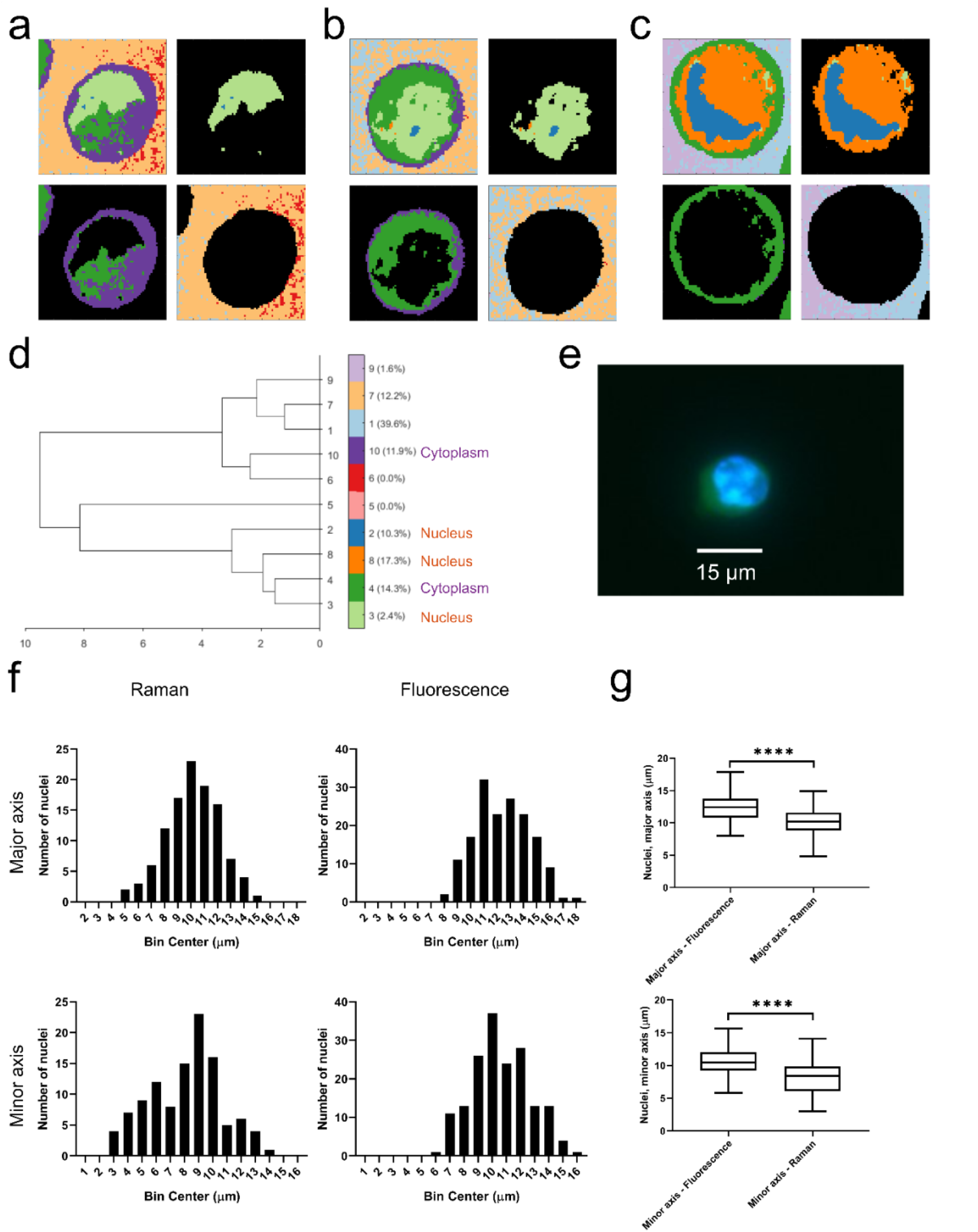
Assessing the cell and nucleus segmentation. **(a-c)** Additional example cell maps (specifically samples 200114-CH12-D4-009 (a), 200212-CH12-D0-002 (b), and 200228-CH12-D0-007 (c)) after common K-means with 10 clusters, as presented in Figure 1a-c. Nucleus (top right), cytoplasm (bottom left), and background (bottom right) associated pixels highlighted. **(d)** Hierarchical clustering analysis plot of the common K-means. The clusters identified as cytoplasm and nucleus are annotated. **(e)** Epifluorescence image of a CH12F3 cell: the nucleus is stained with Hoechst (blue) and the whole cell with SYTO13 (green). **(f)** Histograms showing the nucleus size distribution (major axis, left and minor axis, right) of the cells measured from Raman maps (top) and epifluorescence microscopy images (bottom). **(g)** Quantification of (b). A t-test gave a statistically significant difference between samples (ns.: P>0.05, *: <0.05, **: P<0.01, ***:P<0.001).

### Quantifiable spectral differences between non-activated and activated B cells

CH12F3 CSR in response to CIT treatment was verified by flow cytometry (Figure S2). D0 cells almost exclusively produce IgM BCRs, while a subset of D4 cells have undergone IgM to IgA isotype switching.

PCA analysis was applied to identify the spectral differences between D0 and D4 cells. Four principal components (PC1, PC4, PC5, and PC8) had statistically significant different scores between D0 and D4 cells. Figure S3a-c show the separation between D0 (red) and D4 cells (blue) for these PCs, while Figure S3d-g show their PCA loading spectra.

**Figure S2:**
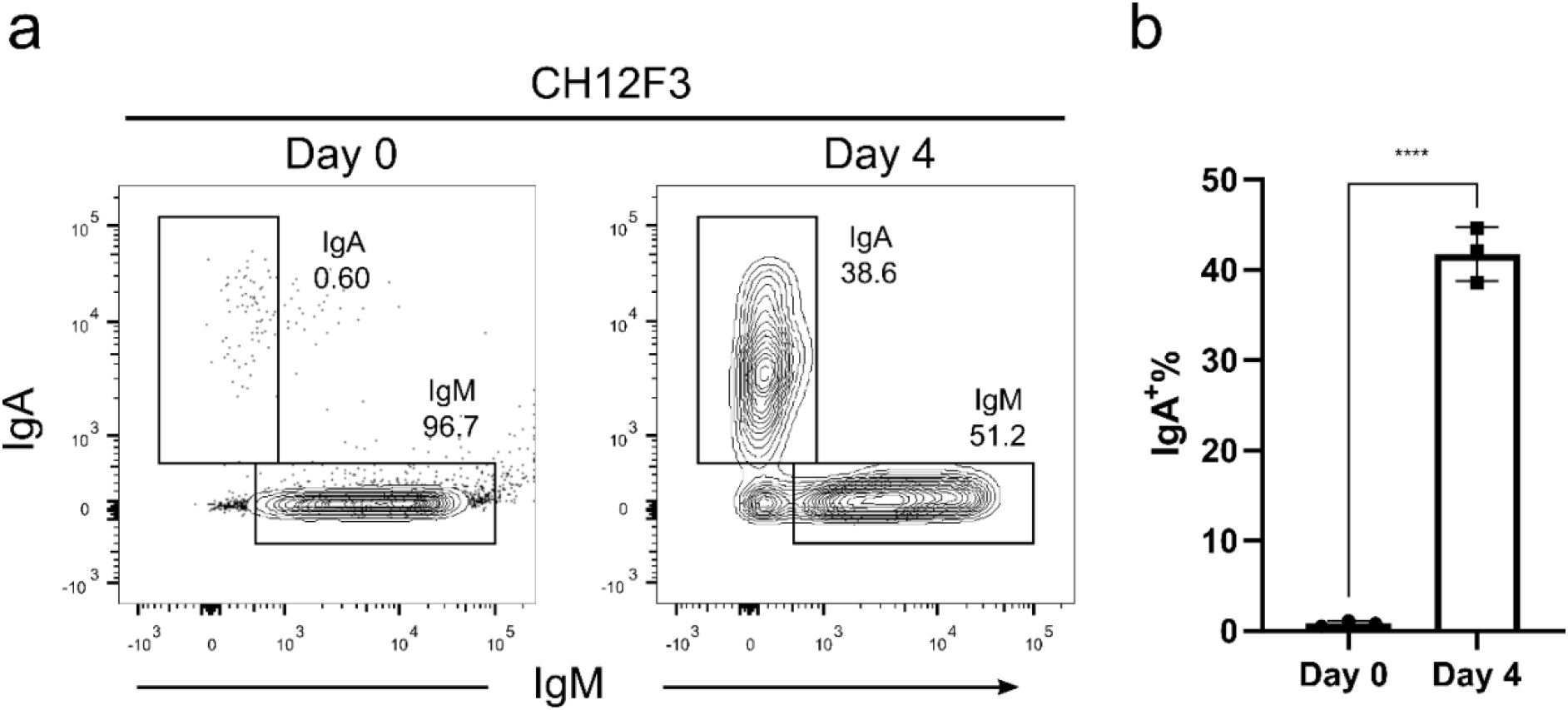
Monitoring CSR in CH12F3 cells. **(a)** CSR in CH12F3 upon CIT stimulation monitored by identifying IgM- and IgA-producing cells using flow cytometry. **(b)** Quantification of IgA+ cells for D0 and D4.

**Figure S3:**
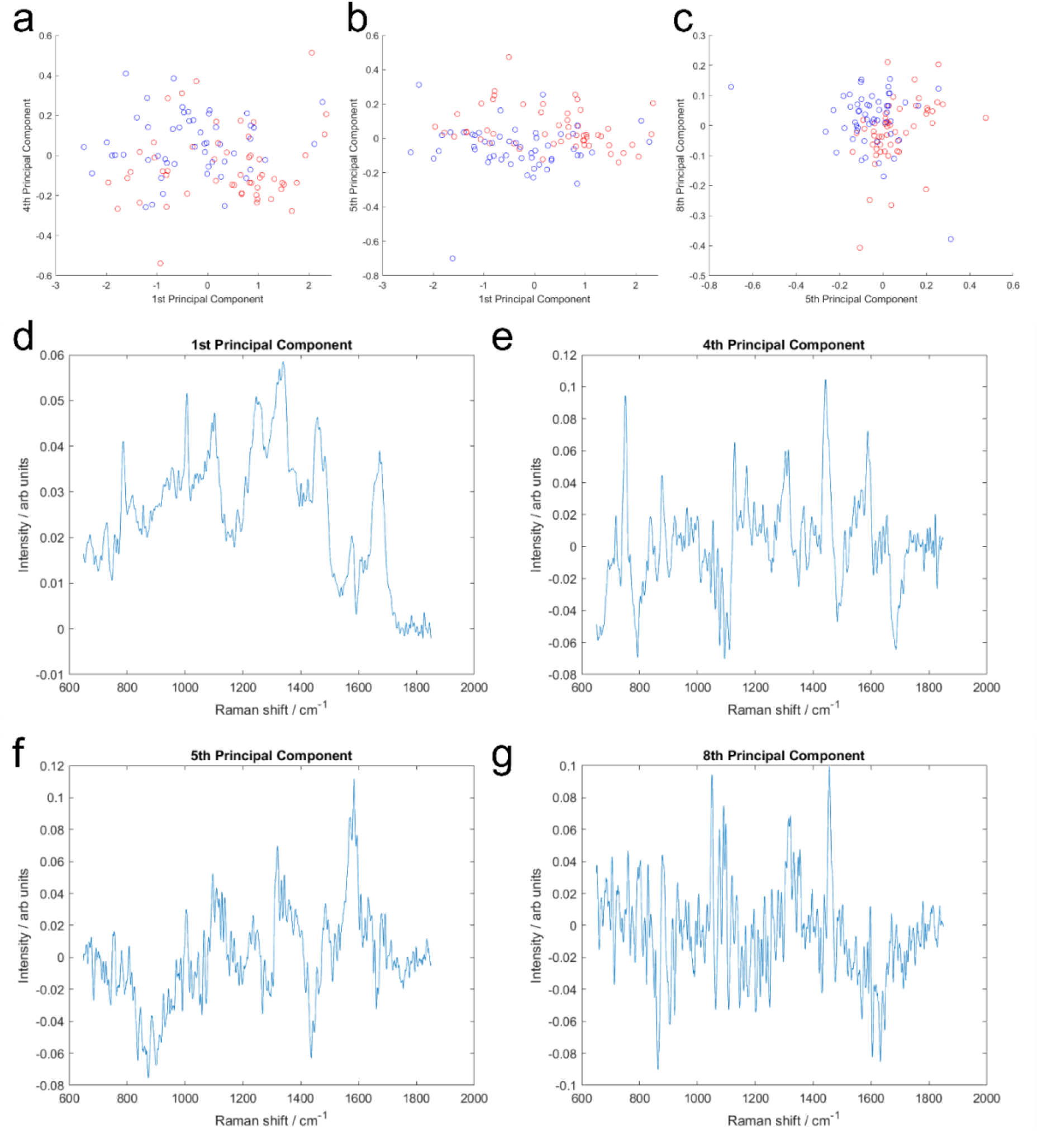
PCA and LDA analysis for discrimination between D0 and D4 cellss. **(a)** PCA scores; PC1 versus PC4. **(b)** PCA scores; PC1 versus PC5. **(c)** PCA scores; PC5 versus PC8. **(d)** PCA loadings; PC1. **(e)** PCA loadings; PC4. **(f)** PCA loadings; PC5. **(g)** PCA loadings; PC8.

### Predicting Raman data from transcriptomic data using PLS regression models

The linear correlation between Raman data and transcriptomic data, as determined by the PLS analysis, is visualised in Figure S4a for component 1. The D0 samples cluster together, as do the D4 samples. This is the basis for the model which enabled the prediction of Raman data from transcriptomic data (Figure 4b-c and Figure S4b-e).

In addition to the Raman nucleus data (Figure 4 and Figure S4), Raman whole cell and cytoplasm data were also used for PLS analysis (Figure S5). The whole cell data analysis results were largely identical to those of the nucleus data analysis (Figure S5a). For the cytoplasm data, the results differed more. However, the linear correlation was still clear (Figure S5b), and it was still possible to accurately predict Raman data from transcriptomic data (Figure S5c-f).

**Figure S4:**
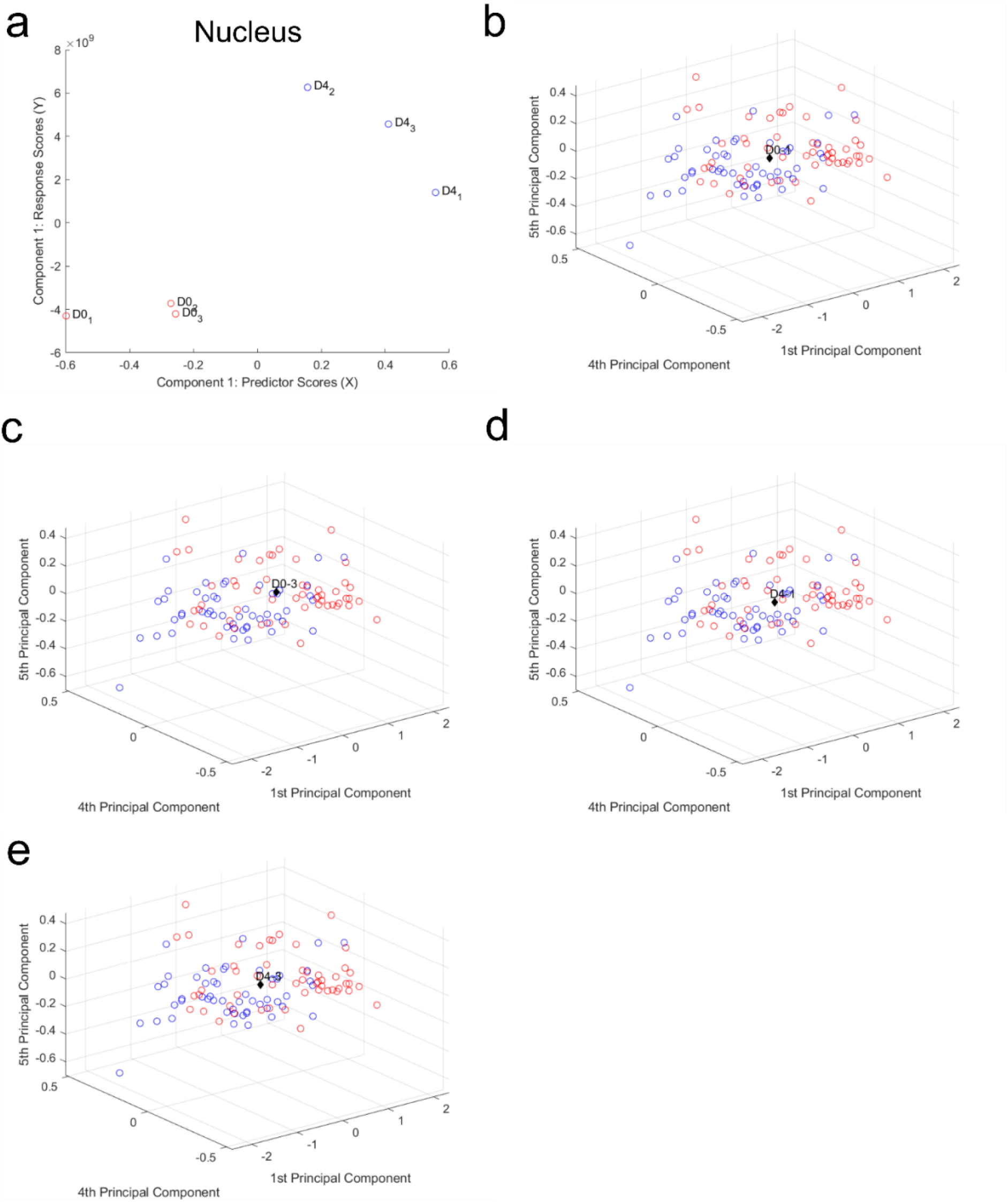
PLS regression model can predict Raman data from transcriptomic data. **(a**) PLS regression analysis shows a linear correlation between Raman nucleus data and transcriptomic data for component 1. **(b-e)** Raman scores predicted from transcriptomic read counts. D0-1 (b), D0-3 (c), D4-1 (d), and D4-3 (e) plotted with the single cell scores from components 1, 4 and 5.

**Figure S5:**
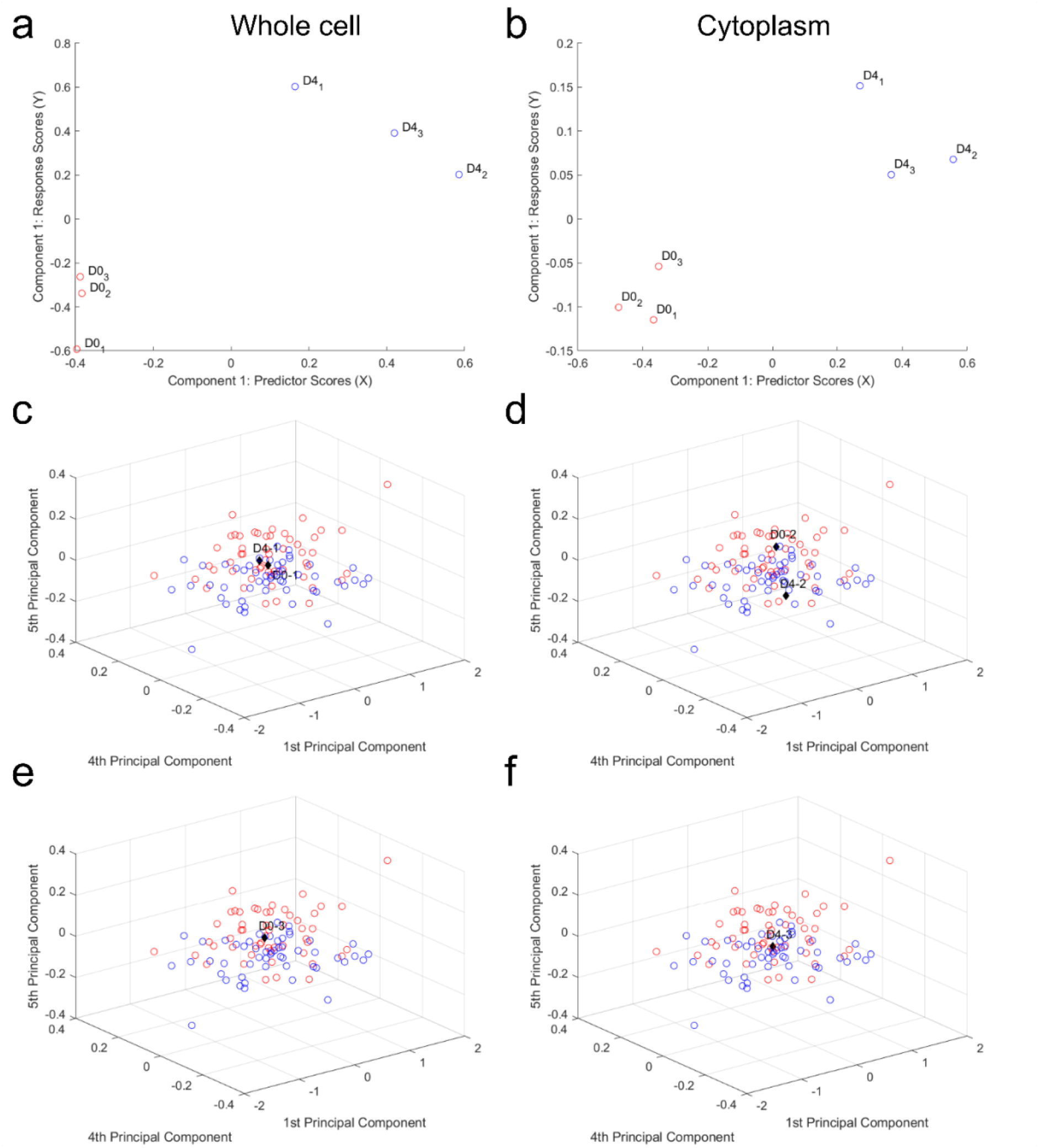
PLS regression model correlates Raman whole cell and cytoplasm data with transcriptomic data. **(a)** PLS regression analysis shows a linear correlation between Raman whole cell data and transcriptomic data for component 1. **(b)** PLS regression analysis shows a linear correlation between Raman cytoplasm data and transcriptomic data for component 1. **(c-f)** Raman cytoplasm PLS model: Raman scores predicted from transcriptomic read counts. D0-1 and D4-1 (c), D0-2 and D4-2 (d), D0-3 (e), D4-3 (f) plotted with the single cell scores from PC1, PC4, and PC5.

**Table S1:** VIP list for Whole cell Raman data PLS regression analysis

| VIP score | Transcript name | Transcript type | Gene name |  |
| --- | --- | --- | --- | --- |
| 41.68549 | ENSMUSG00000064339 | Mt_rRNA | mt-Rnr2 | mitochondrially encoded 16S rRNA |
| 31.63403 | ENSMUSG00000037742 | protein_coding | Eef1a1 | Elongation factor 1-alpha 1 |
| 30.71642 | ENSMUSG00000095079 | IG_C_gene | Igha | immunoglobulin heavy constant alpha |
| 30.42896 | ENSMUSG00000092341 | lncRNA | Malat1 | metastasis associated lung adenocarcinoma transcript 1 |
| 29.63411 | ENSMUSG00000076617 | IG_C_gene | Ighm | immunoglobulin heavy constant mu |
| 28.14179 | ENSMUSG00000029580 | protein_coding | Actb | actin, beta |
| 23.71356 | ENSMUSG00000097971 | lncRNA | Gm26917 | predicted gene, 26917 |
| 23.70888 | ENSMUSG00000065037 | misc_RNA | Rn7sk | RNA, 7SK, nuclear |
| 22.0795 | ENSMUSG00000064351 | protein_coding | mt-Co1 | mitochondrially encoded cytochrome c oxidase I |
| 20.93438 | ENSMUSG00000047139 | protein_coding | Cd24a | CD24a antigen |
| 20.11123 | ENSMUSG00000034994 | protein_coding | Eef2 | eukaryotic translation elongation factor 2 |
| 15.38487 | ENSMUSG00000011179 | protein_coding | Odc1 | ornithine decarboxylase, structural 1 |
| 14.87474 | ENSMUSG00000031779 | protein_coding | Ccl22 | chemokine (C-C motif) ligand 22 |
| 14.05003 | ENSMUSG00000032399 | protein_coding | Rpl4 | 60S ribosomal protein L4 |
| 13.63799 | ENSMUSG00000015656 | protein_coding | Hspa8 | heat shock protein 8 |
| 11.70491 | ENSMUSG00000026864 | protein_coding | Hspa5 | heat shock protein 5 |
| 11.58428 | ENSMUSG00000049775 | protein_coding | Tmsb4x | thymosin, beta 4, X chromosome |
| 11.54382 | ENSMUSG00000024359 | protein_coding | Hspa9 | heat shock protein 9 |
| 10.83353 | ENSMUSG00000057113 | protein_coding | Npm1 | Nucleophosmin |
| 10.81621 | ENSMUSG00000022797 | protein_coding | Tfrc | transferrin receptor |

**Table S2:**
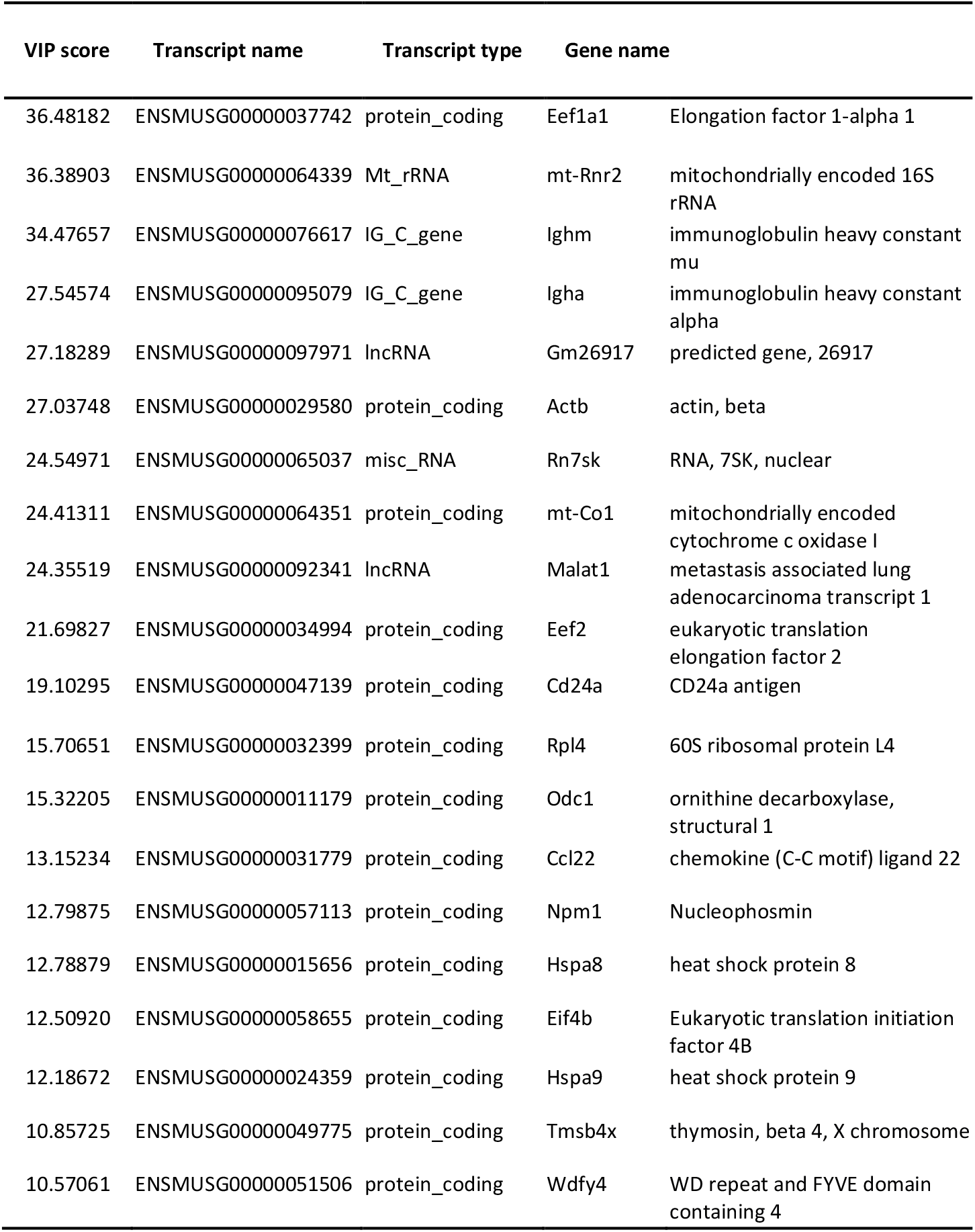
VIP list for Cytoplasm Raman data PLS regression analysis

## References

1. Sanderson MJ, Smith I, Parker I, Bootman MD (2014) Fluorescence Microscopy. Cold Spring Harb Protoc 10:1–36.

2. Levy SE, Myers RM (2016) Advancements in Next-Generation Sequencing. Annu Rev Genomics Hum Genet 17:95–115.

3. Boydston-White S, Gopen T, Houser S, Bargonetti J, Diem M (1999) Infrared spectroscopy of human tissue. V. Infrared spectroscopic studies of myeloid leukemia (ML-1) cells at different phases of the cell cycle. Biospectroscopy 5:219–227.

4. Holman H-YN, Martin MC, McKinney WR (2003) Tracking chemical changes in a live cell: Biomedical applications of SR-FTIR spectromicroscopy. Spectroscopy 17:139–159.

5. Short KW, Carpenter S, Freyer JP, Mourant JR (2005) Raman spectroscopy detects biochemical changes due to proliferation in mammalian cell cultures. Biophys J 88(6):4274–4288.

6. Swain RJ, Jell G, Stevens MM (2008) Non-invasive analysis of cell cycle dynamics in single living cells with Raman micro-spectroscopy. J Cell Biochem 104(4):1427–1438.

7. Morrish RB, et al. (2019) Single Cell Imaging of Nuclear Architecture Changes. Front Cell Dev Biol 7:141.

8. Chen T, et al. (2008) Pharmacodynamic Assessment of Histone Deacetylase Inhibitors: Infrared Vibrational Spectroscopic Imaging of Protein Acetylation Pharmacodynamic Assessment of Histone Deacetylase Inhibitors: Infrared Vibrational Spectroscopic Imaging of Protein Acetylati. Anal Chem 80(16):6390–6396.

9. Zhang F, Huang Q, Yan J, Zhang X, Li J (2015) Assessment of the effect of trichostatin a on hela cells through FT-IR spectroscopy. Anal Chem 87(4):2511–2517.

10. Meade AD, Byrne HJ, Lyng FM (2010) Spectroscopic and chemometric approaches to radiobiological analyses. Mutat Res - Rev Mutat Res 704(1–3):108–114.

11. Lipiec E, et al. (2013) A new approach to studying the effects of ionising radiation on single cells using FTIR synchrotron microspectroscopy. Radiat Phys Chem 93:135–141.

12. Lipiec E, et al. (2014) Monitoring UVR induced damage in single cells and isolated nuclei using SR-FTIR microspectroscopy and 3D confocal Raman imaging. Analyst 139(17):4200–9.

13. Lipiec E, et al. (2014) Synchrotron FTIR shows evidence of DNA damage and lipid accumulation in prostate adenocarcinoma PC-3 cells following proton irradiation. J Mol Struct 1073(C):134–141.

14. Wood B, Tait B, Naughton DMC (2000) Fourier Transform Infrared Spectroscopy as a Method for Monitoring the Molecular Dynamics of Lymphocyte Activation. 54(3):353–359.

15. Wood BR, Tait B, McNaughton D (2000) Fourier-transform infrared spectroscopy as a tool for detecting early lymphocyte activation: A new approach to histocompatibility matching. Hum Immunol 61(12):1307–1315.

16. Ramoji A, et al. (2012) Toward a spectroscopic hemogram: Raman spectroscopic differentiation of the two most abundant leukocytes from peripheral blood. Anal Chem 84(12):5335–5342.

17. Maguire A, et al. (2015) Competitive evaluation of data mining algorithms for use in classification of leukocyte subtypes with Raman microspectroscopy. Analyst 140(7):2473–2481.

18. Brown KL, et al. (2009) Differentiation of alloreactive versus CD3/CD28 stimulated T-lymphocytes using raman spectroscopy: A greater specificity for noninvasive acute renal allograft rejection detection. Cytom Part A 75(11):917–923.

19. Zoladek AB, et al. (2010) Label-free molecular imaging of immunological synapses between dendritic and T cells by Raman micro-spectroscopy. Analyst 135(12):3205–3212.

20. Smith ZJ, Wang J-CE, Quataert SA, Berger AJ (2010) Integrated Raman and angular scattering microscopy reveals chemical and morphological differences between activated and nonactivated CD8+ T lymphocytes. J Biomed Opt 15(3):036021.

21. Pully V V., Lenferink ATM, Otto C (2011) Time-lapse Raman imaging of single live lymphocytes. J Raman Spectrosc 42(2):167–173.

22. Mazur AI, et al. (2013) Vibrational spectroscopic changes of B-lymphocytes upon activation. J Biophotonics 6(1):101–109.

23. Hobro AJ, Kumagai Y, Akira S, Smith NI (2016) Raman spectroscopy as a tool for label-free lymphocyte cell line discrimination. Analyst 141(12):3756–3764.

24. Pavillon N, Hobro AJ, Akira S, Smith NI (2018) Noninvasive detection of macrophage activation with single-cell resolution through machine learning. Proc Natl Acad Sci 115(12):201711872.

25. Ichimura T, et al. (2016) Non-label immune cell state prediction using Raman spectroscopy. Sci Rep 6(1):37562.

26. Germond A, et al. (2018) Raman spectral signature reflects transcriptomic features of antibiotic resistance in Escherichia coli. Commun Biol 1(1). doi:10.1038/s42003-018-0093-8.

27. Kobayashi-Kirschvink KJ, et al. (2018) Linear Regression Links Transcriptomic Data and Cellular Raman Spectra. Cell Syst 7(1):104–117.e4.

28. Smith A, et al. (2018) The culture environment influences both gene regulation and phenotypic heterogeneity in Escherichia coli. Front Microbiol 9(AUG):1–13.

29. Sheppard EC, Morrish RB, Dillon MJ, Leyland R, Chahwan R (2018) Epigenomic modifications mediating antibody maturation. Front Immunol 9(FEB):355.

30. Zheng S, et al. (2015) Non-coding RNA Generated following Lariat Debranching Mediates Targeting of AID to DNA. Cell 161(4):762–773.

31. Vigorito E, et al. (2007) microRNA-155 Regulates the Generation of Immunoglobulin Class-Switched Plasma Cells. Immunity 27(6):847–859.

32. Wang L, Whang N, Wuerffel R, Kenter AL (2006) AID-dependent histone acetylation is detected in immunoglobulin S regions. J Exp Med 203(1):215–26.

33. Stanlie A, Aida M, Muramatsu M, Honjo T, Begum N a (2010) Histone3 lysine4 trimethylation regulated by the facilitates chromatin transcription complex is critical for DNA cleavage in class switch recombination. Proc Natl Acad Sci U S A 107:22190–22195.

34. Ramachandran S, et al. (2016) The SAGA Deubiquitination Module Promotes DNA Repair and Class Switch Recombination through ATM and DNAPK-Mediated gH2AX Formation. Cell Rep 15(7):1554–1565.

35. Xu Z, et al. (2010) 14-3-3 adaptor proteins recruit AID to 5’-AGCT-3’-rich switch regions for class switch recombination. Nat Struct Mol Biol 17(9):1124–35.

36. Pagliara S, Chimerel C, Langford R, Aarts DGAL, Keyser UF (2011) Parallel sub-micrometre channels with different dimensions for laser scattering detection. Lab Chip 11(19):3365–3368.

37. Trapnell C, et al. (2012) Differential gene and transcript expression analysis of RNA-seq experiments with TopHat and Cufflinks. Nat Protoc 7(3):562–578.

38. Metsalu T, Vilo J (2015) ClustVis: A web tool for visualizing clustering of multivariate data using Principal Component Analysis and heatmap. Nucleic Acids Res 43(W1):W566–W570.

39. Zhang Z, Zhang Y, Evans P, Chinwalla A, Taylor D (2017) RNA-seq 2G: Online analysis of differential gene expression with comprehensive options of statistical methods. bioRxiv (6). doi:10.1101/122747.

40. Heng TSP, et al. (2008) The Immunological Genome Project: networks of gene expression in immune cells. Nat Immunol 9(10):1091–1094.

41. Fan Y, et al. (2016) miRNet - dissecting miRNA-target interactions and functional associations through network-based visual analysis. Nucleic Acids Res 44(W1):W135–W141.

42. Li JH, Liu S, Zhou H, Qu LH, Yang JH (2014) StarBase v2.0: Decoding miRNA-ceRNA, miRNA-ncRNA and protein-RNA interaction networks from large-scale CLIP-Seq data. Nucleic Acids Res 42(D1):92–97.

43. Shannon P, et al. (2003) Cytoscape: A Software Environment for Integrated Models of Biomolecular Interaction Networks. (Karp 2001):2498–2504.

44. Ridoux JP, Liquier J, Taillandier E (1988) Raman Spectroscopy of Z-Form Poly[d(A-T)]•Poly[d(A-T)]. Biochemistry 27(2):3874–3878.

45. Kendall C, et al. (2010) Evaluation of Raman probe for oesophageal cancer diagnostics. Analyst 135(12):3038–3041.

46. Stone N, Kendall C, Smith J, Crow P, Barr H (2004) Raman spectroscopy for identification of epithelial cancers. Faraday Discuss 126(1):141–157.

47. Ruiz-Chica AJ, Medina MA, Sánchez-Jiménez F, Ramírez FJ (2004) Characterization by Raman spectroscopy of conformational changes on guanine-cytosine and adenine-thymine oligonucleotides induced by aminooxy analogues of spermidine. J Raman Spectrosc 35(2):93–100.

48. Liu Z, et al. (2008) Circulation and long-term fate of functionalized, biocompatible single-walled carbon nanotubes in mice probed by Raman spectroscopy. Proc Natl Acad Sci U S A 105(5):1410–1415.

49. Huang Z, et al. (2003) Near-infrared Raman spectroscopy for optical diagnosis of lung cancer. Int J Cancer 107(6):1047–1052.

50. Stone N, Kendall C, Shepherd N, Crow P, Barr H (2002) Near-infrared Raman spectroscopy for the classification of epithelial pre-cancers and cancers. J Raman Spectrosc 33(7):564–573.

51. Cheng WT, Liu MT, Liu HN, Lin SY (2005) Micro-Raman spectroscopy used to identify and grade human skin pilomatrixoma. Microsc Res Tech 68(2):75–79.

52. Shetty G, Kendall C, Shepherd N, Stone N, Barr H (2006) Raman spectroscopy: Elucidation of biochemical changes in carcinogenesis of oesophagus. Br J Cancer 94(10):1460–1464.

53. Gniadecka M, Wulf HC, Mortensen NN, Nielsen OF, Christensen DH (1997) Diagnosis of basal cell carcinoma by Raman spectroscopy. J Raman Spectrosc 28(2–3):125–129.

54. Krafft C, Neudert L, Simat T, Salzer R (2005) Near infrared Raman spectra of human brain lipids. Spectrochim Acta - Part A Mol Biomol Spectrosc 61(7):1529–1535.

55. Kamemoto LE, et al. (2010) Near-infrared micro-raman spectroscopy for in vitro detection of cervical cancer. Appl Spectrosc 64(3):255–261.

56. Ribeiro de Almeida C, et al. (2018) RNA Helicase DDX1 Converts RNA G-Quadruplex Structures into R-Loops to Promote IgH Class Switch Recombination. Mol Cell 70(4):650–662.e8.

57. Huong LT, et al. (2013) In Vivo Analysis of Aicda Gene Regulation: A Critical Balance between Upstream Enhancers and Intronic Silencers Governs Appropriate Expression. PLoS One 8(4):1–12.

58. Dedeoglu F, Horwitz B, Chaudhuri J, Alt FW, Geha RS (2004) Induction of activation-induced cytidine deaminase gene expression by IL-4 and CD40 ligation is dependent on STAT6 and NFκB. Int Immunol 16(3):395–404.

59. Delgado-Benito V, et al. (2018) The Chromatin Reader ZMYND8 Regulates Igh Enhancers to Promote Immunoglobulin Class Switch Recombination. Mol Cell 72(4):636–649.e8.

60. Warde-Farley D, et al. (2010) The GeneMANIA prediction server: Biological network integration for gene prioritization and predicting gene function. Nucleic Acids Res 38(SUPPL. 2):214–220.

61. Orthwein A, et al. (2010) Regulation of activation-induced deaminase stability and antibody gene diversification by Hsp90. J Exp Med 207(12):2751–2765.

62. Li J, et al. (2019) The coordination between B cell receptor signaling and the actin cytoskeleton during B cell activation. Front Immunol 10(JAN):1–13.

63. Song W, Liu C, Upadhyaya A (2014) The pivotal position of the actin cytoskeleton in the initiation and regulation of B cell receptor activation. Biochim Biophys Acta - Biomembr 1838(2):569–578.

64. Baarlink C, et al. (2017) A transient pool of nuclear F-actin at mitotic exit controls chromatin organization. Nat Cell Biol 19(12):1389–1399.

65. Hurst V, Shimada K, Gasser SM (2019) Nuclear Actin and Actin-Binding Proteins in DNA Repair. Trends Cell Biol 29(6):462–476.

66. Caridi CP, Plessner M, Grosse R, Chiolo I (2019) Nuclear actin filaments in DNA repair dynamics. Nat Cell Biol 21(9):1068–1077.

67. Benevides JM, Thomas GJ (1983) Characterization of DNA structures by Raman spectroscopy: High-salt and low-salt forms of double helical poly(dG-dC) in H2O and D2O solutions and application to B, Z and A-DNA. Nucleic Acids Res 11(16):5747–5761.

68. Ridoux JP, Liquier J, Taillandier E (1988) Raman Spectroscopy of Z-Form Poly[d(A-T)]•Poly[d(A-T)]. Biochemistry 27:3874–3878.

69. Wittig B, Dorbic T, Rich A (1991) Transcription is associated with Z-DNA formation in metabolically active permeabilized mammalian cell nuclei. Proc Natl Acad Sci U S A 88(6):2259–2263.

70. Shin SI, et al. (2016) Z-DNA-forming sites identified by ChIP-Seq are associated with actively transcribed regions in the human genome. DNA Res 23(5):477–486.

